# How bumblebees coordinate path integration and body orientation at the start of their first learning flight

**DOI:** 10.1101/2022.11.04.515210

**Authors:** TS Collett, T Robert, E Frasnelli, A. Philippides, N Hempel de Ibarra

## Abstract

The start of a bumblebee’s first learning flight from its nest provides an opportunity to examine the bee’s learning behaviour on its initial view of the nest’s unfamiliar surroundings. Bumblebees like many other ants, bees and wasps learn views of their nest surroundings while facing their nest. A bumblebee’s first fixation of the nest is a coordinated manoeuvre in which the insect faces the nest with its body oriented towards a particular visual feature within its surroundings. The manoeuvre’s utility is that during return flights after foraging bees, when close to the nest, adopt the same preferred body-orientation (Hempel de Ibarra et al., 2009; Robert et al., 2018). A translational scan oriented orthogonally to the bee’s body-orientation helps the bee reach the preferred conjunction of nest-fixation and body-orientation.

How does a bee, unacquainted with its surroundings, know when it is facing its nest? The details of nest-fixation argue that, like desert ants (Fleischmann et al., 2018), the bee relies on path integration. Path integration gives bees continuously updated information about the current direction of their nest and enables them to fixate the nest when the body points in the appropriate direction. We relate the three components of the coordinated manoeuvre to events in the central complex, noting that nest fixation is in egocentric coordinates, whereas body orientation and flight direction within the visual surroundings of the nest are in geocentric coordinates.

## Introduction

When ants, bees, or wasps of many species first leave their nest they perform, in its immediate vicinity, what have come to be called ‘learning walks’ for ants and ‘learning flights’ for bees and wasps (reviews: Collett and Zeil, 2018; Zeil and Fleischmann, 2019; Zeil et al., 1996). The insects’ learning behaviour consists of a directed path during which an insect periodically looks back to face its nest. Nest facing is crucial as it is then that an insect most probably learns the visual surroundings of the nest (Bates, 1863; Lehrer, 1993). In some bees (Hempel de Ibarra et al., 2009) and wasps (Zeil, 1993a) the insect faces its nest while pointing in a preferred compass direction (North in the case of the bumblebee *Bombus terrestris*) or at a favoured visual feature in its surroundings. A view recorded when facing the nest enables a returning insect to use that memory to guide its return to the nest (Dewar et al., 2014, Hempel de Ibarra et al., 2009; Robert et al., 2018; Zeil, 1993b).

In ants, the amount of nest facing depends on the environment in which a species lives (Fleischmann et al., 2017). Nest facing lasts longer when the nest is located within vegetation than it does for species with nests in bare surroundings, suggesting that the duration of nest facing depends on what must be learnt about visual features near the nest. Bumblebees tend to nest in holes in the ground that are often abandoned by other animals and are quite commonly in undergrowth. Correspondingly, bumblebees tend to fixate the nest for relatively long periods. The Results start with an analysis of these fixations.

How do insects leaving their nest with no knowledge of the landscape around the nest manage to face it? A definitive answer has come from desert ants. By applying an artificial magnetic field to ants engaged in learning walks in their natural surroundings, it was shown that ants rely on path integration (Fleischmann et al., 2018). Path integration holds a vector of an insect’s current direction and distance to the nest throughout a learning walk or flight. It would thus enable an ant to face the nest at will. When the direction of the field, was shifted, the ants responded by facing towards their nest in the perceived magnetic direction, ignoring other compass cues, demonstrating thereby that the direction of the nest was available through path integration.

Because colonies of *Bombus terrestris* are available commercially with new workers emerging from their pupae, we can be sure about a bee’s visual experience and know that the bees we test have not seen the visual surroundings of the nest before their learning flight. The bumblebees’ lack of visual experience means that facing the nest is, as in ants, most likely implemented through path integration. This notion is supported by examining the precision with which they face the nest. Next, we ask whether the bees, while they fixate the nest, face in a common direction relative to their newly viewed surroundings.

Since the answer to this question was yes, we looked for correlations between the bees’ preferred body orientation when they first face the nest and the direction of their return flights to the nest. Lastly, we examine details of the manoeuvre that leads to nest facing.

## Materials and Methods

Experiments were conducted in a greenhouse (8×12 m floor area) at the Streatham campus of the University of Exeter, UK. Bumblebees, *Bombus terrestris audax* (Linnaeus 1758), from commercially reared colonies (Koppert, Haverhill, UK), were marked individually with coloured number tags. The colony was placed under a table, the ‘nest table’, and we recorded the flights of worker bees as we allowed them to leave their nest, one at a time, through a hole in the centre of the table. Three black cylinders (17×5 cm) were placed in a 120° arc around the nest hole with their centres 24.5 cm from the hole. Another table with a feeder and the same arrangement of cylinders was placed 5m from the nest table. Both tables were covered with white gravel.

The behaviour of bees leaving the nest and in some cases on their returns after feeding was recorded continuously during the experiments at 50 frames s^−1^ with video cameras (Panasonic HC-V720, HD 1080p) that were hung 1.35 m above each table and captured an area of about 70×90 cm in an image of 1920×1080 pixels.

### Experimental Procedure

A bee colony, kept indoors overnight, was taken in the morning to the greenhouse. The box containing the colony was placed beneath the nest table with the nest-box connected to a hole in the centre of the table via a series of tubes used to control the exits and re-entrances of individual bumblebees from the nest

All the bees contributing to this study were recorded on the first occasion that they left their nest. At the end of their learning flight from the nest they flew around the greenhouse before being caught and placed on the feeder. Recordings of flights come from Robert et al. (2018) (N=18 bees) and from an unpublished study (N=16 bees) that followed the same procedures.

### Data analysis and statistics

All videos were examined and clipped with video-editing software (Adobe CS6). We discarded the few flights in which bees landed during the learning flight or flew directly away from the nest. Bumblebee learning flights have two phases: an initial phase in which bees fly close to the nest and are low on the ground and a second phase in which they gain height and fly further from the nest (Collett et al., 2013; Linander et al., 2018; Lobecke et al., 2018; Philippides et al., 2013; Riabinina et al., 2014). We stopped analysis of the flights once the bees had travelled 5cm from the nest. The positions and body orientations of the bees were extracted from the video recordings using custom-written code in MATLAB which allows corrections by hand ensuring orientations are accurate to ∼5° (details in Hempel de Ibarra, 2009). We analysed the bee’s body orientation relative to the nest and also its body orientation relative to its surroundings. For the latter, we chose one of the landmarks as a reference point.

A single bout of nest facing was mostly quite short, but it could last for almost a second. We use the term ‘fixation’ for a bout of nest facing in which the bee faced the centre of the nest with a precision of ±10° for at least 4 frames (i.e. 80ms). The 80ms minimum duration of a fixation comes from the plateaus and saccades seen in bumblebee head movements (Riabinina et al., 2014). If just a single frame within the fixation lay outside the limit of ±10°, the frame was included as part of the fixation. To work with separate fixations, we only included those fixations that had an interval of at least 200ms between the end of one fixation and the start of the next one. An example of an exclusion is given in Fig 1.

**Figure 1.**
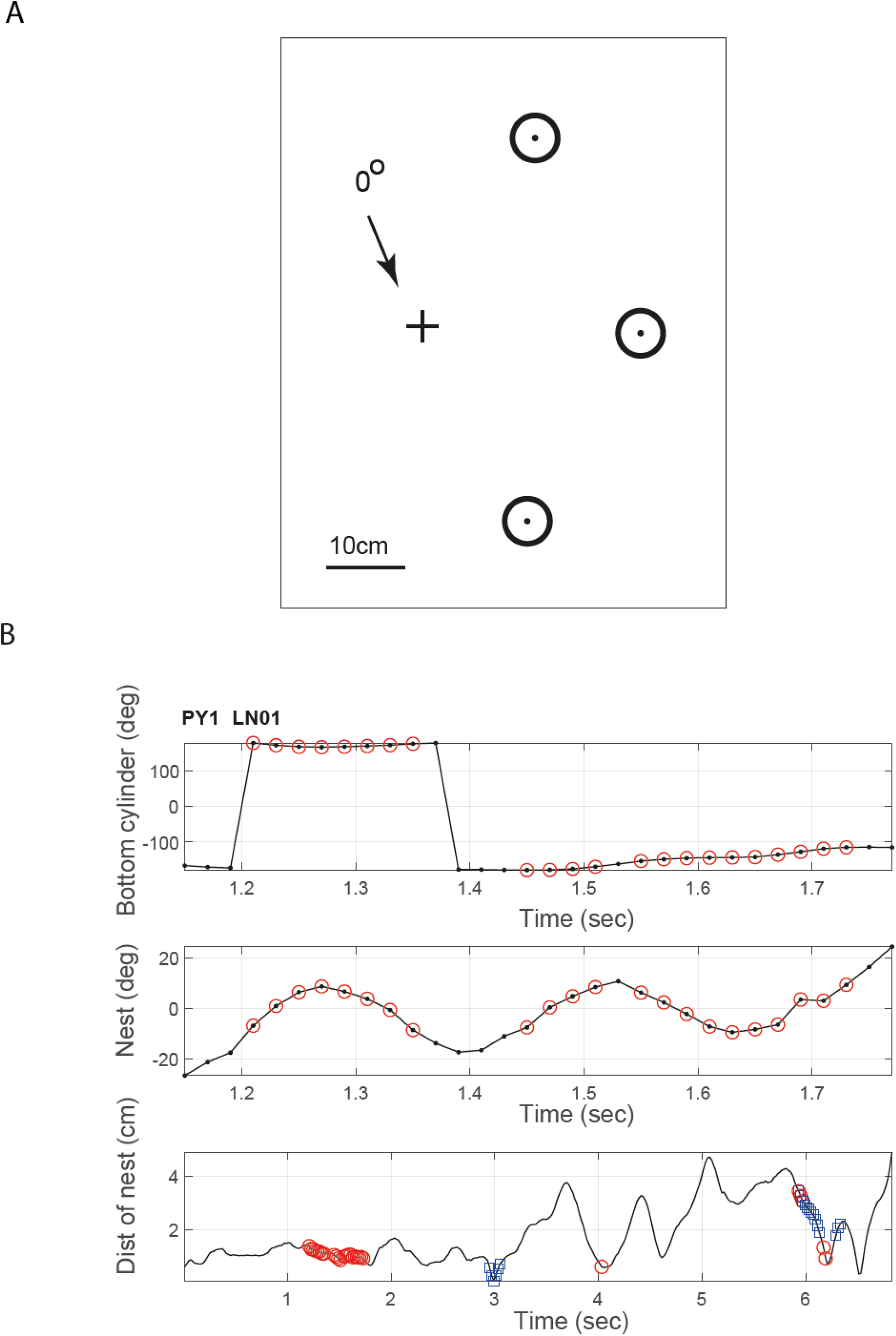
**A**. Layout of cylinders (circles) around the nest (+). The reference direction (0°) is parallel to the line between the nest and the bottom cylinder. **B**. An example of a fixation set within the first 5cm of a learning flight. Each box plots a parameter of a single bee’s first learning flight from the nest over a section of the flight and shows how fixations are selected. Top: Body orientation relative to bottom cylinder (0°). To make fixations independent of each other, we only selected fixations that are separated by at least 200ms. Red circles are frames in which bee faced the nest (±10°). Because the two putative fixations are separated by only 80ms the longer of the two fixations (in this case the second one) is retained and the other rejected. Middle: Body orientation relative to nest. The single frame outside the fixation at ca. 1.52s is accepted as part of the fixation. Note its proximity to 10°. The longer period of frames outside the fixation at 1.4s is excluded Bottom: Whole flight showing the bee’s distance from nest, Blue is when bee faces the bottom cylinder. In this case, the first nest fixation does not coincide with the bee facing the bottom cylinder. The two do coincide in the last nest fixation. Thus, the first fixation would not have been included in the histogram of Fig. 4C, but the last fixation would contribute to the histogram of Fig 4D.

To determine whether fixations were longer than one might expect by chance, we performed a Monte Carlo analysis with 100,000 repetitions. For each repetition, the frames of all the learning flights over the analysed distance (0 to 5cm) were randomised. Fixations were then extracted from all the randomised frames. Statistical tests on the data were performed in R and in Matlab using the CircStat tool box (Berens, 2009).

## Results

### Nest facing at the start of a learning flight

Bumblebees during the start of their first learning flights spend most of the time facing in the general direction of the nest. Translational speeds, plotted against body angles relative to the nest decline sharply close to zero, but rotational speeds have a more ragged decline around zero. The numbers at the top of the plot give the number of frames in the associated bins, indicating the predominance of nest facing., as occurs more generally in bumblebee learning flights (Hempel de Ibarra et al., 2009).

Although earlier papers (Hempel et al. 2009; Philippides at al., 2013) excluded the initial part of the flight when bees were very close to the nest and accumulated data over the whole flight, the picture presented there was similar, with a predominance of nest facing and a decrease in translational speed when that occurred.

### Fixations

The bees tend to face the nest in bouts of consecutive frames that we term fixations (see Methods). Their durations have a broad spread with a maximum close to a second (Fig. 2D). We looked for ways to show that these durations are longer than would be expected by chance, given the general prevalence of nest facing. A first attempt was to examine fixations of points perpendicular to the nest and test whether they were shorter than nest-fixations. Although there were fewer of these fixations and none were longer than 20 frames (Fig. 2 E, F), a comparison of the distribution of 57 perpendicular fixations with the distribution of nest-fixations found no difference in fixation lengths (Mann-Whitney U-test: z-score = -0.25, p= 0.40).

**Figure 2.**
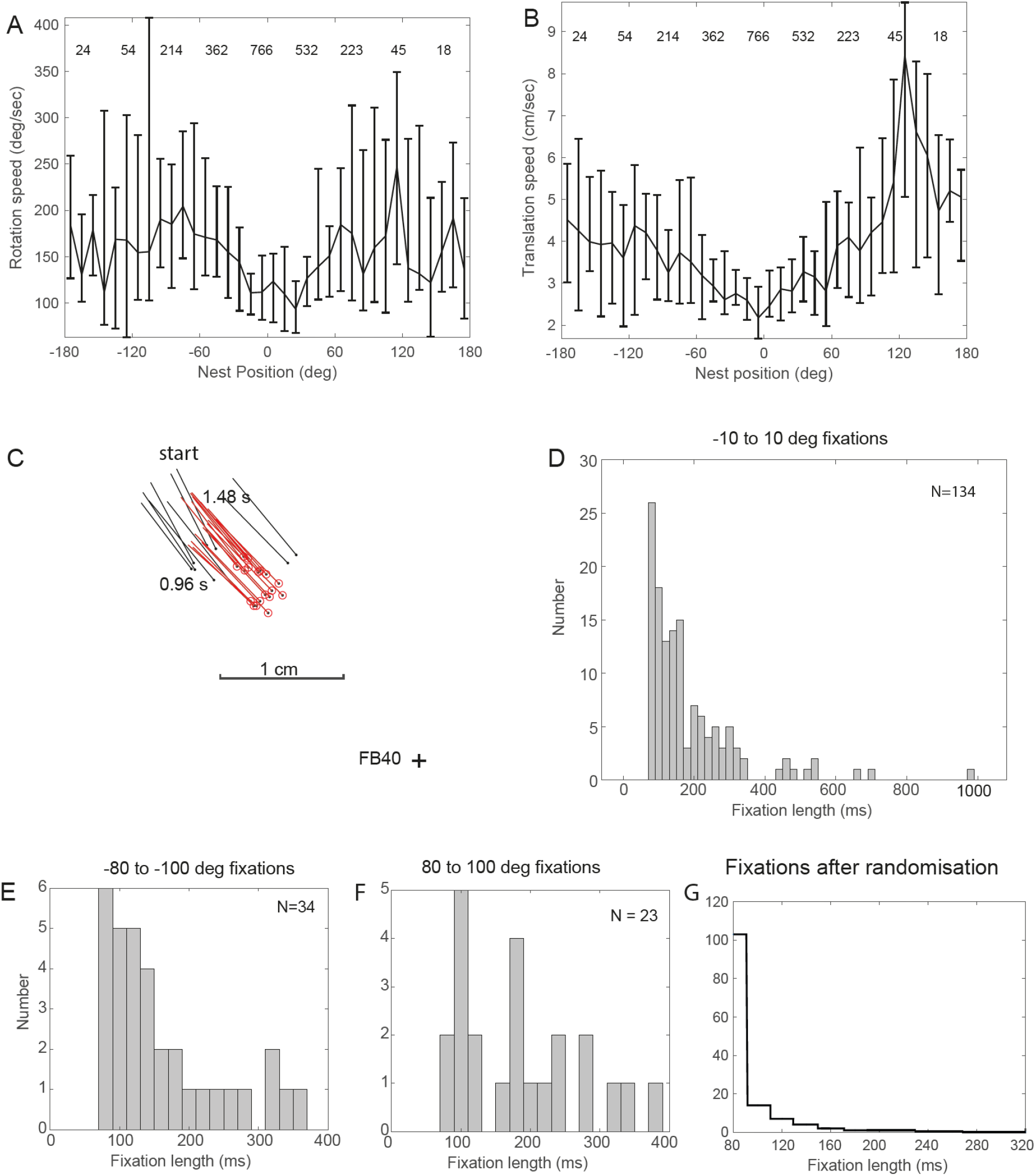
Nest-fixations. **A** and **B**. Median rotational and translational speeds of 33 bees during their first learning flights as a function of body orientation relative to the nest. Data is accumulated within 10° bins with the speed range indicated by bars from the 25^th^ to 75^th^ percentiles of the data in each bin. Numbers above the plot show how many frames the bees spent in that bin and emphasise the preponderance of nest facing. Error bars are interquartile range. **C**. Example of nest-fixation. Ball and stick give the position of the head and the orientation of bee FB40’s fixation of the nest every 20ms. Red is the fixation. To show this bee’s precision, the limits here are ±5°. Points outside this limit are black. + is placed here and in other figures at the central point of the nest hole. Times give the start and end of the fixation. **D**. The durations of all nest-fixations of 32 bees in msec. **E** and **F**. Distributions of fixation durations when bees face towards the two points that are perpendicular to the nest. **G**. Distribution of the durations of nest-fixations after the data from the learning flights were randomised.

The next step was to examine the length of fixations when the data were randomised in a Monte Carlo analysis with 100,000 repetitions (see Data analysis in Materials and Methods). Fixations extracted from all the randomised frames were pooled and a plot of the frequency of their relative lengths (Fig 2G) shows a sharp drop in the frequency of fixations longer than 5 frames. A comparison (Fig. 3) between the real and randomised distributions of fixation lengths, assuming independence between bins, establishes that the real fixations are significantly longer (Mann-Whitney U test: z-score = -2.10106, p= 0.01786). The same is true for fixations extracted using the same process that was adopted for the randomised results, but before randomisation (Mann-Whitney U test: z-score = -1.91162, p=0.02807). We conclude that nest-fixations are part of the bee’s intended behaviour, justifying our focus on fixations in the following sections.

**Figure 3.**
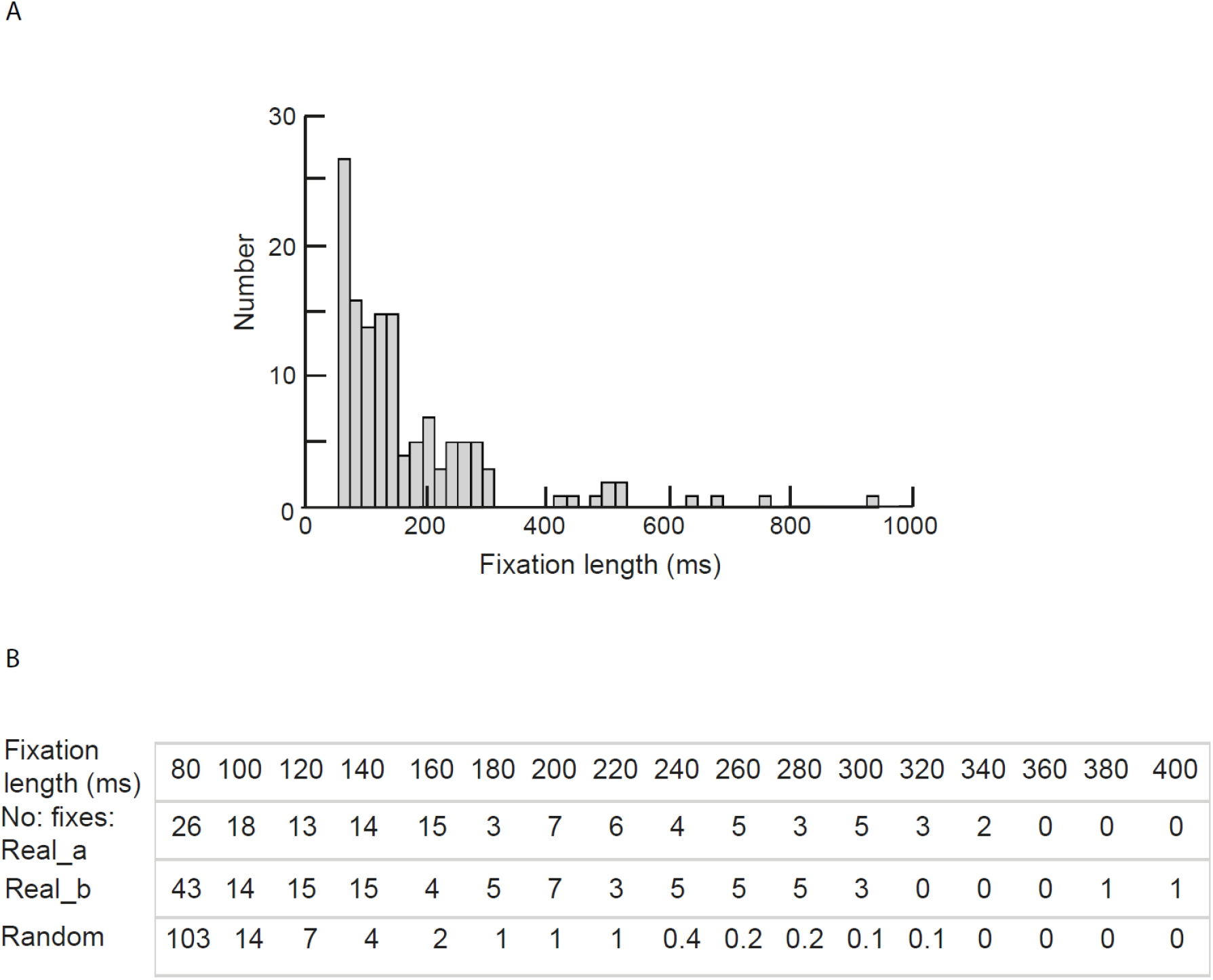
**A**. Length of fixations extracted for Monte Carlo analysis from 32 learning flights before randomisation. Plot is very similar to the corresponding plot extracted manually (Fig. 2D), the major difference being that criteria of 200 ms separation was not applied. **B**. Testing whether real fixations are longer than those occurring if all the frames from all the analysed nest-learning flights are randomised 100,000 times. All rows are normalised to a total of 134 fixations. The analysis stops when fixation duration reaches 400ms and the frequency of further random fixations is well below 1. The diagram has four rows. The top row shows the length of the fixations from 80 to 400 ms. The next three rows give the number of occurrences of each fixation length. First row: fixation lengths as shown in Fig 2D; second row: fixation lengths as shown in in Fig. 3A; third row: fixation lengths after randomisation. Compared with both rows of real fixations, randomisation leads to more short fixations and fewer long fixations.

### Path integration and fixations

What tells bees that are unacquainted with their surroundings to lower their rotational and translational speeds during nest-fixations? At the very start of the flight bees move away and then turn back to fixate the nest. They are thus unlikely to recognise the nest entrance and to guide their turn towards the nest using visual cues. Visual recognition is particularly unlikely because fixations occur when bees are close to the nest entrance (median distance = 2.1 cm, IQR = 1.7 cm, Fig. 4A). For some learning flights the nest was marked by a 5cm purple ring (Robert et al., 2018), so at 2cm height it subtends ca 100° on the ventral region of the eye. In other flights, the outer diameter of the exit tube was 2.2cm (Fig. 4B) so it would subtend ca. 60°. A ±10° angle of fixation at 2 cm means that bees can pinpoint the nest centre to within ca 0.7cm (Fig. 4A). A reliance on path integration (Honkanen et al., 2019; Mittelstaedt and Mittelstaedt, 1980) thus seems more plausible. Throughout the flight, path integration holds a vector of an insect’s current direction and distance to the nest enabling the bee to face the nest at will.

**Figure 4.**
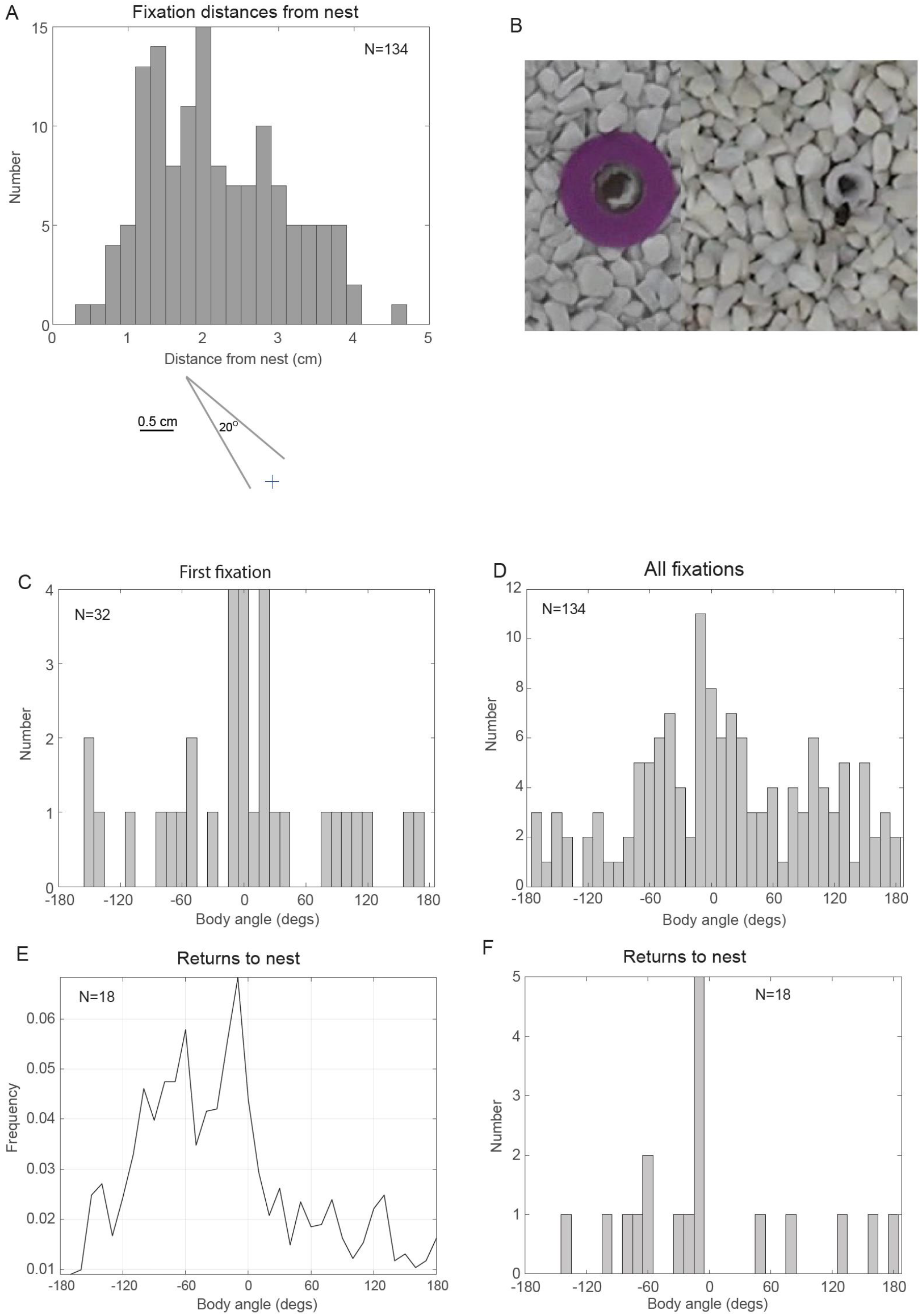
Body orientation and distance from the nest during nest-fixations and body orientation during return flights. **A**. Horizontal distances of mid-point of fixations from the nest. Diagram below illustrates the span covered by a fixation at 2cm from the nest. **B**. Photo showing nests with and without purple ring. Ring diameter is 5cm and outer tube 2.2cm. **C, D**. Mean of bees’ body orientation relative to the bottom cylinder during their first (C) and all fixations (D). **E,F**. Body orientation during return flights when bee is within 10 cm of nest. E. Frequency of body orientations accumulated over all bees. F. Peak orientation of each of the 18 bees.

### Preferred body orientations during nest-fixations and on return flights

We took the bees’ body orientation during their fixations of the nest as their orientation relative to the direction of the line from the nest to the bottom cylinder (Fig. 4B) at the midpoint of each fixation. Peak body orientation was 0°. The effect is especially clear on the first fixation of the flight (cf. Fig. 4C and Fig. 4D). We used the peak value of the body orientation of each bee for statistics, giving a sample size of 32 for the data in both Fig. 4C and D. Circular statistics applied to these distributions gave a circular mean of -3.7°, R=0.39 for the first fixations and 11.6°, R=0.25 for all fixations. The V test with a prediction of 0° indicates that body orientations are not distributed randomly around 360° (first fixations eval=12.54, p<0.001; all fixations eval=24.11, p<0.001).

For 18 of the bees from Robert et al. (2018) there were matching return flights to the nest after the bees’ first visit to a flower. We extracted the bees’ body orientations during their return flight, starting once they were within 10 cm of the nest. Over this last section of the return, the distribution of body angles peaks at -10°, just to the left of the bottom cylinder, but with a wide scatter (Fig. 4E).

The peak and the scatter are reflected in the bees’ individual performance (Fig. 4F). For each bee we took the peak value of its body orientation during its return. Five bees had a peak at -10°. The V test with a prediction of 0° was significant (eval=5.54, p<0.04), R value = 0.38, circular mean = 36.74. This behaviour is consistent with earlier data that the bees’ preferred body orientations during learning flights is replicated on return flights (Hempel de Ibarra et al., 2009; Robert et al., 2018). The new analysis adds that the conjunction between facing the nest during fixations and facing the bottom cylinder is clearest close to the start of the first learning flight, which raises the question whether the first fixations might be longer than later fixations, giving a greater chance of facing the bottom cylinder. But a comparison of the lengths of the first fixations with those of all the subsequent fixations indicates no difference (Mann-Whitney U-test: z-score = -0.88193. p= 0 .37886).

The correlation between the bees’ body orientation during learning and return flights coupled with the bees’ tendency during learning flights to face the bottom cylinder when facing the nest gives the returning bee a simple strategy for reaching the nest. The bee maintains its body orientation while on the lookout for a view that matches its memory.

### Reaching a coincidence between nest-fixation and preferred body orientation

We noticed that just before bees faced both the nest and the bottom cylinder, they performed a translational scan that was roughly perpendicular to the line between the nest and the bottom cylinder (Fig. 5A). The data are limited. 11 of the 32 analysed bees exhibited a clear overlap between facing the nest and the bottom cylinder during their first fixation of the nest. In 5 cases, the body reached the preferred body orientation before nest facing occurred (e.g., bee FB22, Fig.5A). In 4 cases the order was reversed. Twice bees faced the nest and the bottom cylinder at the same time (e.g. bee FG3, Fig. 5A). Further examples of the flight trajectories are given in Fig. 5B. If body orientation is close to zero during the scan, this arrangement enables the bee to stop scanning when path integration tells it that its preferred body orientation angle coincides with the direction of the nest. Conversely, facing the nest (Fig. 4A) while scanning means that the bee can stop scanning when it reaches its preferred body orientation.

**Figure 5.**
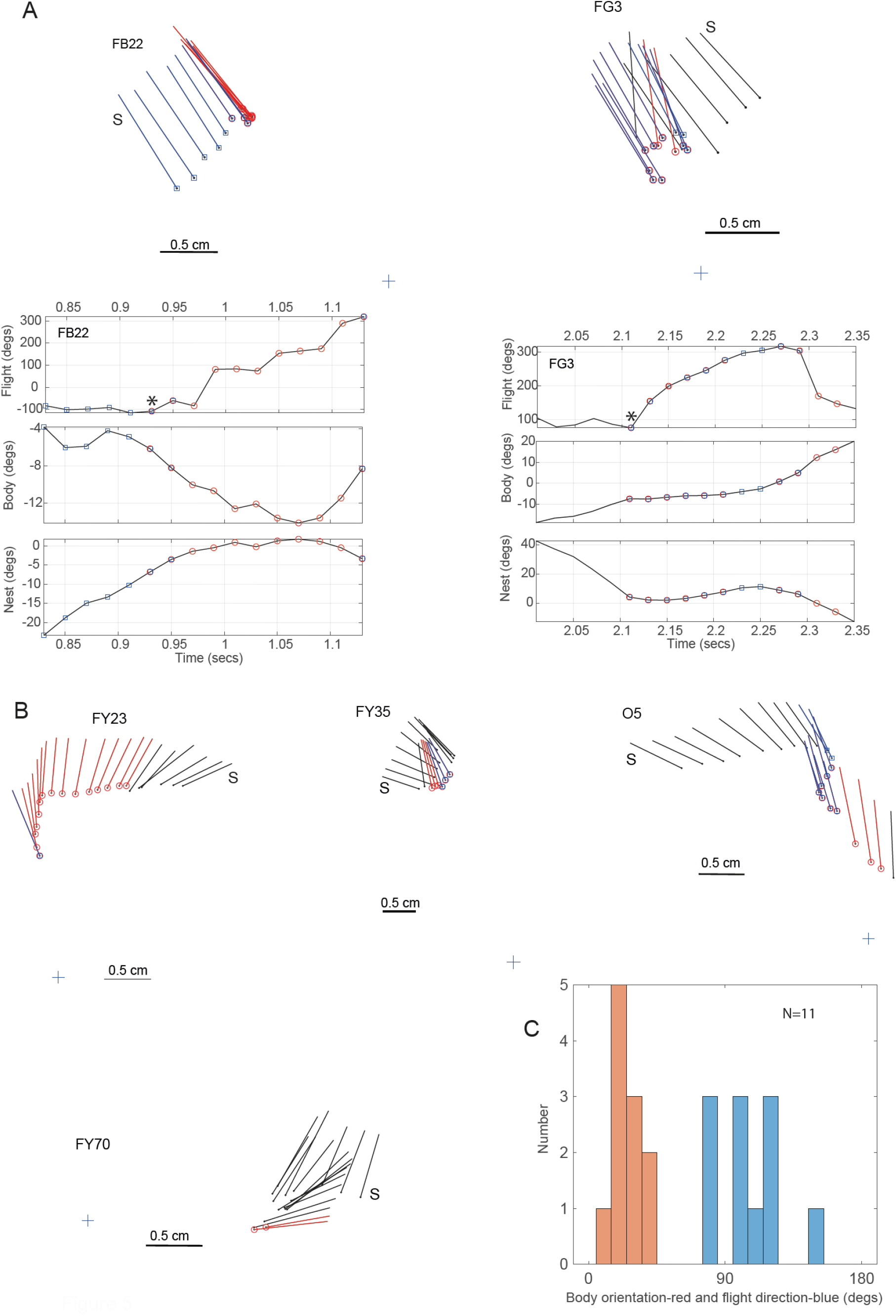
Translational scans prior to the coincidence of simultaneously facing the nest and the bottom cylinder. **A**. Flight paths of the scans and fixations of two bees. S indicates the start of the scan. Time plots of the scans of bees FB22 and FG3 give, from top to bottom, the bees’ flight direction, their body orientation relative to line from nest to bottom cylinder, and lastly their body orientation relative to the nest. Nest facing frames (±10°) are red and bottom cylinder facing frames (±10°) blue. * marks start of coincidence. **B**. The scans and nest-fixations of four more bees. **C**. Histogram (bottom right) plots absolute values of body orientation and flight direction relative to line from nest to bottom cylinder at the midpoint of the scans of the 11 bees.

To quantify flight direction and body orientations during the scan, their values were extracted at the midpoint of each scan. With the signs of the angles ignored, circular statistics on the distributions of flight directions and body orientations (Fig 5C) show that the two distributions differ. For flight directions, the V test with a prediction of 90° was significant (eval=10.21, p<0.0001), R value = 0.94, circular mean = 107.17°; but with a prediction of 0° the values were not significant (eval= -3.06, p=0.90). In contrast, with the V test applied to body orientation the relations reversed, a prediction of 0° was significant (eval=9.84, p<0.0001), R value = 0.99, circular mean = 25.17°. But with a prediction of 90° the values were at chance (eval= -0.27, p= 0.55).

## Discussion

The data here add to what we already know about learning flights in *Bombus terrestris* by showing what happens at the start of a bee’s first learning flight, when it is unfamiliar with the visual surroundings outside its nest. Despite this ignorance, bees during their first fixation of the nest tend to face the point in their surroundings that corresponds to the body orientation that will be adopted by bees returning to their nest.

We also have indications that, like ants (Fleischmann et al., 2018), nest facing is accomplished with the aid of path integration. Lastly, we have learnt that the bee controls its flight direction during a scan of the scene. This scan helps the bee reach a conjunction between nest facing and its preferred viewing point in the nest surroundings.

### Flight parameters and the central complex in the brain

A bumblebee’s body orientation within its surroundings, its fixation of its nest, and its flight direction contribute in an organised way to the bee’s ability to memorise views during learning flights. These three parameters of its flight are most likely controlled by the central complex. In *Drosophila*, the direction in which a fly faces is encoded spatially within the ring-like ellipsoid body (Seelig and Jayaraman, 2015). The ring consists of 8 tiles with each sensitive to a particular direction. Only a single tile is active at any time and the fly points in the direction encoded by the active tile. Visual input to the ring is carried by a population of ring neurons with inhibitory processes that connect all the other tiles. In consequence, the active ring cell determines which point in the scene attracts the fly’s attention (Fisher, 2019). This mechanism is well suited to enable a bee to face the bottom cylinder.

Nest-fixations most likely involve path integration and much of the circuitry supporting path integration resides within the fan-shaped body of the central complex (Stone et al., 2017). A further actor, flight direction, is needed to generate the bee’s scan. In *Drosophila*, flight direction, independently of body direction, is also computed within the fan-shaped body (Lu et al., 2022, Lyu et al. 2022). Our data suggest that the three flight parameters are closely coordinated in generating the scan that precedes a nest-fixation (Figs. 5, 6a). How the central complex might implement this coordination is not yet clear.

**Figure 6.**
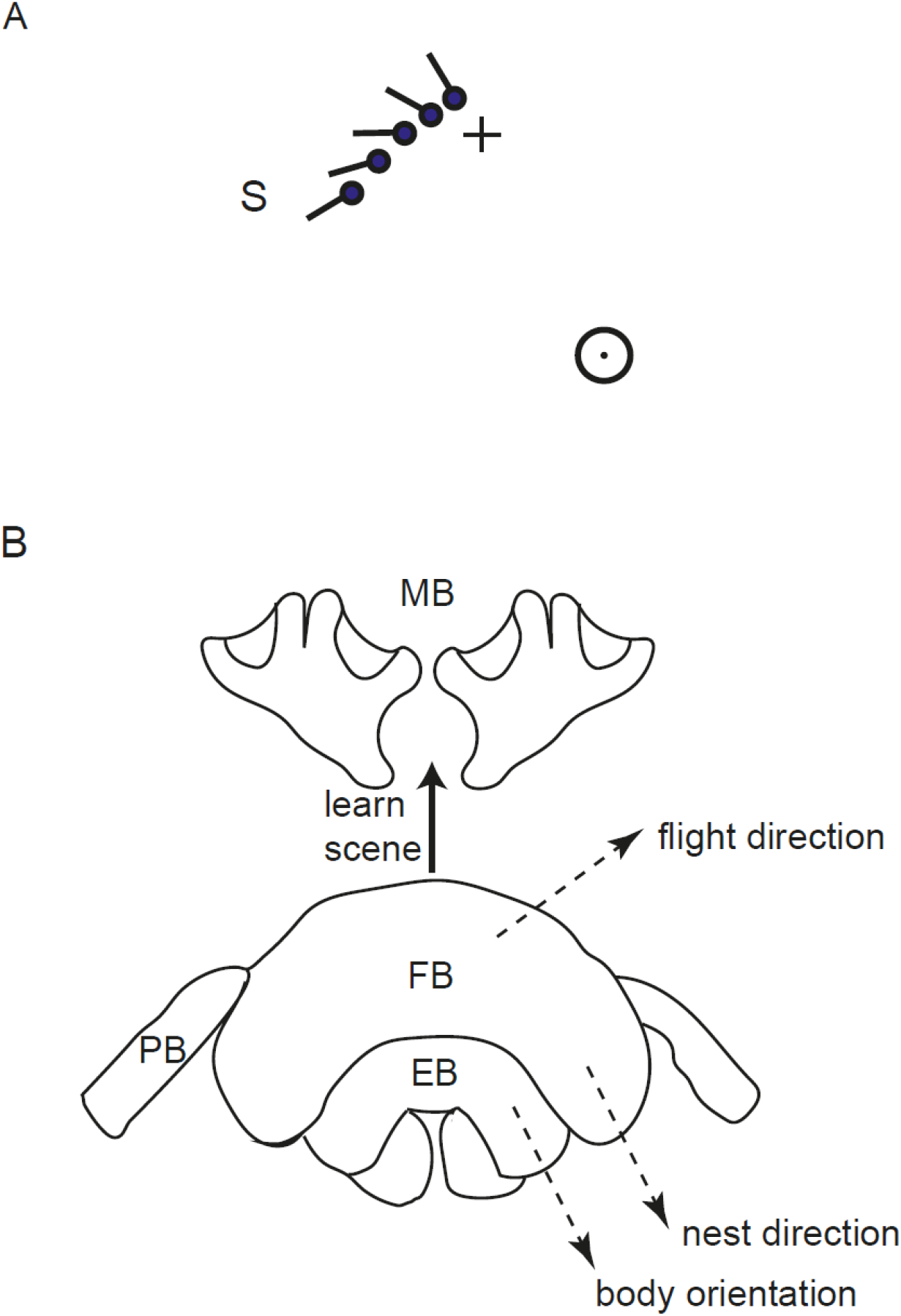
Translational scan and sketch of central complex and mushroom body with suggested interactions between the two structures. **A**. Sketch of the way that a translational scan perpendicular to the line between the nest and bottom cylinder can aid the conjunction of nest facing and the bee’s preferred facing direction - a time when bees may memorise the scene around the nest. Stick and ball signify the orientation and head of the bee. S is start of scan. Figure shows a scan that might occur if the nest is fixated before the conjunction. Rotation would be slower for scans in which the bottom cylinder is fixated first. **B**. MB-mushroom body, EB-ellipsoid body, PB-protocerebral bridge, FB-fan-shaped body. Learning is likely to occur in MB when body orientation is 0° as encoded in EB and the bee at the same time faces the nest as computed through path integration in FB. Flight direction is also elaborated in FB.

In contrast, it is known that the views obtained through the central complex when the bee faces the nest in its preferred body orientation are probably stored within the mushroom body (Buehlmann et al., 2020; Kamhi et al., 2020; Strausfeld, 2012). An apposite connection between the two structures is indicated in *Drosophila* by a relation between walking direction and dopamine activity within the mushroom body (Zolin et al., 2021). Such a partition of tasks between the central complex and the mushroom body (Fig. 6) suggests that if the input from the mushroom body to the central complex were to be experimentally interrupted a bee may be able perform a normal learning flight, but without learning the scene in a useful way.

## Funding

The research was supported by a research grant (RPG-2012-677) to NHI and TSC, and an Emeritus Followship (EM-2016-066) to TSC, both from the Leverhulme Trust. AP was funded by an EPSRC grant

## Data and code availability statement

The research data supporting this publication are openly available from the University of Exeter’s institutional repository at: https://doi.org/XXX/exe.XXX (to be uploaded).

## Notes

### Competing Interest Statement

The authors have declared no competing interest.

## References

Bates, H. W. (1863). The naturalist on the river Amazons. Reprinted 1969. London, UK: JM Dent and Sons.

Berens, P. (2009). CircStat: a MATLAB toolbox for circular statistics. J. Stat. Software 31, 1–21.

Buehlmann, C., Wozniak, B., Goulard, R., Webb, B., Graham, P. and Niven, J. E. (2020). Mushroom bodies are required for learned visual navigation, but not for innate visual behavior, in ants. Curr. Biol. 30, 3438-3443. e2.

Collett, T. S., Hempel de Ibarra, N., Riabinina, O. and Philippides, A. (2013). Coordinating compass-based and nest-based flight directions during bumblebee learning and return flights. J. Exp. Biol. 216, 1105–13.

Collett, T. S. and Zeil, J. (2018). Insect learning flights and walks. Curr. Biol. 28, R984–R988.

Dewar, A. D., Philippides, A., & Graham, P. (2014). What is the relationship between visual environment and the form of ant learning-walks? An in silico investigation of insect navigation. Adapt.Behav. 22, 163–179.

Fleischmann, P. N., Grob, R., Müller, V. L., Wehner, R. and Rössler, W. (2018). The geomagnetic field is a compass cue in Cataglyphis ant navigation. Curr. Biol. 28, 1440-1444. e2.

Fleischmann, P. N., Grob, R., Wehner, R. and Rössler, W. (2017). Species-specific differences in the fine structure of learning walk elements in Cataglyphis ants. J. Exp. Biol. 220, 2426–2435.

Hempel de Ibarra, N., Philippides, A., Riabinina, O. and Collett, T. S. (2009). Preferred viewing directions of bumblebees (Bombus terrestris L.) when learning and approaching their nest site. J. Exp. Biol. 212, 3193–3204.

Honkanen, A., Adden, A., da Silva Freitas, J. and Heinze, S. (2019). The insect central complex and the neural basis of navigational strategies. J. Exp. Biol. 222, jeb188854.

Kamhi, J. F., Barron, A. B. and Narendra, A. (2020). Vertical lobes of the mushroom bodies are essential for view-based navigation in Australian Myrmecia ants. Curr. Biol. 30, 3432-3437. e3.

Lehrer, M. (1993). Why do bees turn back and look? J. Comp. Physiol. A 172, 549–563.

Linander, N., Dacke, M., Baird, E. and de Ibarra, N. H. (2018). The role of spatial texture in visual control of bumblebee learning flights. J. Comp. Physiol. A 204, 737–745.

Lobecke, A., Kern, R. and Egelhaaf, M. (2018). Taking a goal-centred dynamic snapshot as a possibility for local homing in initially naïve bumblebees. J. Exp. Biol. 221, jeb168674.

Lu, J., Behbahani, A. H., Hamburg, L., Westeinde, E. A., Dawson, P. M., Lyu, C., Maimon, G., Dickinson, M. H., Druckmann, S. and Wilson, R. I. (2022). Transforming representations of movement from body-to world-centric space. Nature 601, 98–104.

Lyu, C., Abbott, L. F., & Maimon, G. (2022). Building an allocentric travelling direction signal via vector computation. Nature, 601, 92–97.

Mittelstaedt, M. L. and Mittelstaedt, H. (1980). Homing by path integration in a mammal. Naturwiss. 67, 566–567.

Philippides, A., Hempel de Ibarra, N., Riabinina, O. and Collett, T. S. (2013). Bumblebee calligraphy: the design and control of flight motifs in the learning and return flights of Bombus terrestris. J. Exp. Biol. 216, 1093–104.

Riabinina, O., Hempel de Ibarra, N., Philippides, A. and Collett, T. S. (2014). Head movements and the optic flow generated during the learning flights of bumblebees. J. Exp. Biol. 217, 2633–2642.

Robert, T., Frasnelli, E., Hempel de Ibarra, N. and Collett, T. S. (2018). Variations on a theme: bumblebee learning flights from the nest and from flowers. J. Exp. Biol. 221, jeb172601.

Seelig, J. D. and Jayaraman, V. (2015). Neural dynamics for landmark orientation and angular path integration. Nature 521, 186–191.

Stone, T., Webb, B., Adden, A., Weddig, N. B., Honkanen, A., Templin, R., Wcislo, W., Scimeca, L., Warrant, E. and Heinze, S. (2017). An anatomically constrained model for path integration in the bee brain. Curr. Biol. 27, 3069-3085. e11.

Strausfeld, N. J. (2012). Arthropod brains: evolution, functional elegance, and historical significance: Belknap Press.

Zeil, J. (1993a). Orientation flights of solitary wasps (Cerceris sphecidae, Hymenoptera) I. Description of flight. J. Comp. Physiol. A 172, 189–205.

Zeil, J. (1993b). Orientation flights of solitary wasps (Cerceris; sphecidae; hymenoptera). II: Similarities between orientation and return flights and the use of motion parallax. J. Comp. Physiol. A 172, 207–222.

Zeil, J. and Fleischmann, P. N. (2019). The learning walks of ants (Hymenoptera: Formicidae). Myrmecological News 29, 93–110.

Zeil, J., Kelber, A. and Voss, R. (1996). Structure and function of learning flights in ground-nesting bees and wasps. J. Exp. Biol. 199, 245–252.

Zolin, A., Cohn, R., Pang, R., Siliciano, A. F., Fairhall, A. L. and Ruta, V. (2021). Context-dependent representations of movement in Drosophila dopaminergic reinforcement pathways. Nature Neurosci. 24(11), 1555-1566.

